# HiDRA-seq: High-Throughput SARS-CoV-2 Detection by RNA Barcoding and Amplicon Sequencing

**DOI:** 10.1101/2020.06.02.130484

**Authors:** Emilio Yángüez, Griffin White, Susanne Kreutzer, Lennart Opitz, Lucy Poveda, Timothy Sykes, Maria Domenica Moccia, Catharine Aquino, Ralph Schlapbach

**Author notes:** These authors contributed equally to this work (Emilio Yángüez, Griffin White).

## Abstract

The recent outbreak of a new coronavirus that causes a Severe Acute Respiratory Syndrome in humans (SARS-CoV-2) has developed into a global pandemic with over 6 million reported cases and more than 375,000 deaths worldwide. Many countries have faced a shortage of diagnostic kits as well as a lack of infrastructure to perform necessary testing. Due to these limiting factors, only patients showing symptoms indicating infection were subjected to testing, whilst asymptomatic individuals, who are widely believed to be responsible for the fast dispersion of the virus, were largely omitted from the testing regimes. The inability to implement high throughput diagnostic and contact tracing strategies has forced many countries to institute lockdowns with severe economic and social consequences. The World Health Organization (WHO) has encouraged affected countries to increase testing capabilities to identify new cases, allow for a well-controlled lifting of lockdown measures, and prepare for future outbreaks. Here, we propose HiDRA-seq, a rapidly implementable, high throughput, and scalable solution that uses NGS lab infrastructure and reagents for population-scale SARS-CoV-2 testing. This method is based on the use of indexed oligo-dT primers to generate barcoded cDNA from a large number of patient samples. From this, highly multiplexed NGS libraries are prepared targeting SARS-CoV-2 specific regions and sequenced. The low amount of sequencing data required for diagnosis allows the combination of thousands of samples in a sequencing run, while reducing the cost to approximately 2 CHF/EUR/USD per RNA sample. Here, we describe in detail the first version of the protocol, which can be further improved in the future to increase its sensitivity and to identify other respiratory viruses or analyze individual genetic features associated with disease progression.

## Introduction

Over 350,000 deaths worldwide have resulted from complications resulting from Severe Acute Respiratory Syndrome Coronavirus 2 (SARS-CoV-2) infection with another 6 million reported infected by this virus, causing a major pandemic. The overall public health and economic burden of this pandemic has yet to be realized and will only become apparent in the coming years. Switzerland alone has recorded one of the highest numbers of COVID-19 (the disease caused by caused by SARS-CoV-2) cases per capita in the world^1^. Thus, efficient and sensitive detection assays of SARS-CoV-2 are essential in managing this pandemic, as evidence suggests that the virus is most contagious on or before symptom onset. Furthermore, asymptomatic cases and the so-called “super spreaders” have broadly contributed to the dissemination of the virus, as reports from South Korea suggest^2^. Comprehensive contact tracing, tracking the spread of viral transmission, clearly requires an increase of mass testing to a population scale.

The vast majority of the currently available SARS-CoV-2 diagnostic assays are based on the amplification of specific loci in the viral genome through Real-Time Quantitative Polymerase Chain Reaction (RT-qPCR). RT-qPCR assays have been the gold standard in clinical diagnostics due to their high sensitivity. However, this outbreak has demonstrated that there exists a general lack of infrastructure for such population-scale testing, in addition to a limited supply of reagents for RT-qPCR tests. Furthermore, the outcome of RT-qPCR is a binary result (positive or negative) for the loci interrogated. The main drawback being the lack of genotypic information in RT-qPCR assays, which could enable the mapping of the spread and transmission, as well as the monitoring of the evolution of the etiological agent, which is crucial for vaccine development.

NGS has revolutionized biomedical research in the last 15 years and is increasingly impacting clinical diagnostics and the practice of medicine. Our aim was to develop a diagnostic assay that could profit from the power and sensitivity of these technologies, not only for viral detection but also for viral classification. In this study, we present HiDRA-seq, a low-cost, high-throughput targeted approach for SARS-CoV-2 infection diagnosis and for potentially tracing outbreak origin and tracking transmission. In recent months, several protocols have been developed for SARS-CoV-2 diagnosis using NGS. However, most of the proposed methods are based on the amplification of the entire viral genome, which is time consuming, rather expensive, and require specific kits for enrichment (amplicon or hybridization based) as well as a comprehensive de novo pipeline to analyze the data and advanced bioinformatic pipelines^3^. We propose a midway and rapidly implementable option that uses genomics lab infrastructure and reagents available in large NGS facilities, bypassing the need for commercially available kits and the limitations in the global chain of production. HiDRA-seq combines reverse transcription using barcoded oligo-dT primers, adapted from mcSCRB-seq^4^, with the addition of virus-specific amplicon generation and sequencing. The protocol targets a region of the putative ORF10, which is highly conserved in the different SARS-CoV-2 isolates sequenced to date. The targeted region is located near the 3’-end of the genome^5,6^, so HiDRA-seq can capture both the viral genomic RNA (gRNA) as well as all subgenomic RNA transcripts (sgRNA) generated in infected cells.

We would like to emphasize that our approach consists of a viral enrichment, followed by the generation of a small amount of short read sequencing data and the use of a basic bioinformatics pipeline for downstream mapping and diagnosis. The small amount of short read sequencing data required to correctly diagnose an individual allows the multiplexing of hundreds to thousands of patients in one sequencing run and is an affordable reality at a price of approximately 2 CHF/EUR/USD per sample (from extracted RNA to diagnosis). Furthermore, HiDRA-seq can be adapted by a wide variety of short read sequencers with diverse outputs across the clinics. The nature of the enrichment step in this protocol offers an enormous versatility, as it can be tailored to any other respiratory virus or organism of interest that produces poly-adenylated transcripts. The implementation of a rapid and versatile approach such as HiDRA-seq would definitely enable a more efficient outbreak management.

## Results

### Protocol description

Since the beginning of the SARS-CoV-2 pandemic, various research groups have been working on the development of alternative testing protocols to bypass the shortage of standard diagnostic kits and enable population scale testing. With this idea, we have developed HiDRA-seq, a high throughput, rapidly implementable solution that uses standard lab infrastructure and reagents in medium sized NGS facilities (**Figure 1A**). The reverse transcription and cDNA pooling strategies are adapted from the mcSCBR-seq protocol^4^. Using a low volume liquid handling robot, the patients’ RNA (previously extracted from swab samples) is distributed in 384-well plates containing indexed oligo-dT primers. A short reverse transcription is performed to generate barcoded cDNA and the contents of each well are pooled into a single tube. Following bead purification and exonuclease treatment to digest the excess of unbound primers, the pooled cDNA is used as template in a PCR reaction with a forward primer specific to SARS-CoV-2 and a reverse primer binding to a sequence incorporated with the oligo-dT in the reverse transcription. Both primers contain the sequences required to generate an Illumina sequencer compatible library in a final PCR reaction, in which the amplicon pool can be barcoded, which allows the multiplexing of multiple 384-well plates in a single sequencing run. After sequencing, the reads are de-multiplexed based on both the plate and the sample barcodes and used in an automated analysis pipeline to distinguish samples containing SARS-CoV-2-derived reads, which are diagnosed as positive.

**Figure 1.**
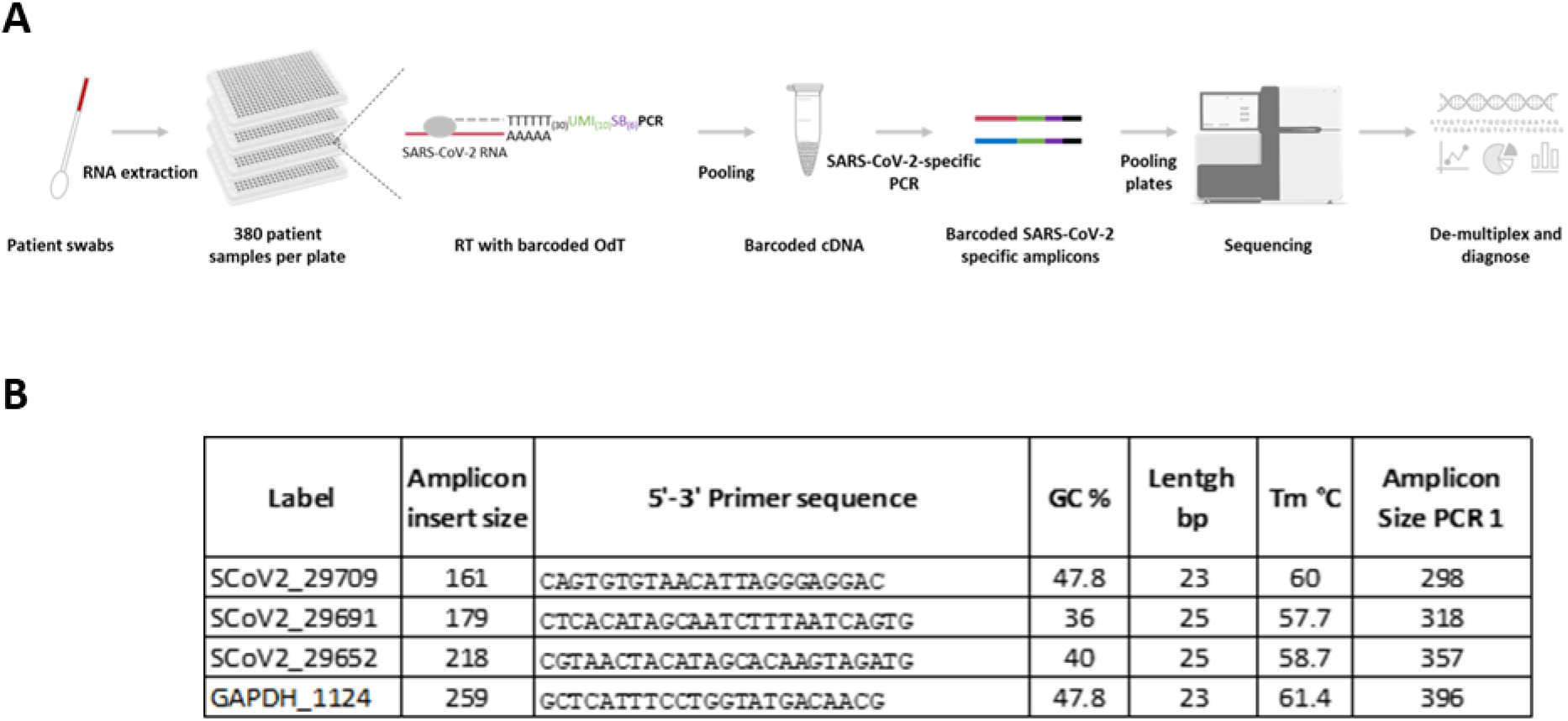
(A) Schematic representation of the protocol. Using a low volume liquid handling robot, the patients’ RNA is distributed in 384-well plates containing indexed oligo-dT primers. Barcoded cDNA is generated by reverse transcription and pooled into a single tube. Libraries are produced from SARS-CoV-2-specific amplicons and sequenced. Reads are de-multiplexed based on both a plate and a sample barcode and used for downstream diagnostic analysis. **(B) Amplicon primer design.** The table contains the sequences of the forward primers used to generate the different amplicons. The reverse primer is common to all amplicons and it anneals to the 3’-end of the barcoded oligo-dT primers used to generate cDNA in the reverse transcription. The primer labels contain the position of the first sequenced base. Insert length was calculated using the start of the poly-A sequence. SARS-CoV-2 NC_045512_2 sequence and GAPDH NM_002046.4 sequence were used as reference. For the theoretical size of the amplicon after PCR 1 the Nextera Tag (34bp), the length of the barcoded Oligo-dT-primer (80bp) and the primer sequence was added.

### SARS-CoV-2-specific amplicon design

We designed three different partially over-lapping amplicons in order to compare their performance and, simultaneously, simulate the combination of three different patient plates that need to be de-multiplexed upon sequencing. For the amplicon design, highly conserved regions between SARS-CoV-2 isolates were identified by creating a whole genome alignment of the European SARS-CoV-2 sequences (n = 1435, sequence identity = 98.96%). The identified sequences are not conserved in other human coronavirus. As cDNA is barcoded using indexed oligo-dT, the forward primers must be located close to the poly-A sequence in the viral genome to obtain amplicons of a reasonable size. Moreover, primers binding to the 3’ of the genome can efficiently capture both the viral genomic RNA (gRNA) as well as all subgenomic RNA transcripts (sgRNAs) generated in infected cells, which may increase the sensitivity of the method. With these premises, we identified a 100% conserved region in the putative ORF10 of SARS-CoV-2. The GC content in this region is highly variable, so primers were designed to target sequences of lower GC contents (<50%). We designed three forward primers targeting this region that are used to generate three SARS-CoV-2 specific amplicons (**Figure 1B**). The three amplicons were tested separately in order to compare their performance. For the human internal control, we followed a similar strategy and designed a forward primer located in close proximity to the poly-A sequence of human GAPDH (Figure 1B). This ensures that a fragment that can be sequenced is always generated even in SARS-CoV-2 negative samples. The reverse primer is common to all amplicons and it anneals to the 3’-end of the barcoded oligo-dT used to generate the cDNA in the reverse transcription.

### Initial data quality control

For the implementation of the protocol, 91 anonymized clinical samples containing RNA extracted from patients’ swaps were kindly provided by Dr. Michael Huber and Dr. Jürg Böni (Institute of Medical Virology, UZH, Switzerland). These clinical samples were transferred to four identical quadrants in a 384-well plate, such that each sample would be processed in quadruplicate for each of the three SARS-CoV-2 amplicons designed and tested for the study. The samples were processed using a Mosquito HV liquid pipetting robot (SPT Labtech), as indicated in materials and methods, and sequenced in both an Illumina MiniSeq (R1=16 bp, i7=8 bp, R2=50 bp) and NovaSeq6000 sequencers (data not shown, R1=16 bp, i7=8 bp, R2=150 bp). The 8 bases barcode was used, in this case, to discriminate the different amplicons but it could alternatively be used to de-multiplex several patient plates. The first 6 bases in read one were used to demultiplex the reads generated from the different patient samples. UMI correction was not applied in this version of the protocol. The 50 bases in read 2 were used to distinguish between positive and negative patients by mapping reads against the SARS-CoV-2 genome. After demultiplexing and prior to mapping, the dataset was subjected to standard quality control checks and data filtering. As the input for the protocols was not normalized, a wide distribution in the number of reads per sample was observed (**Figure 2A**). Reads were filtered by abundance per sample/amplicon combination (n < 250 for one sample/amplicon combination). 250 reads per sample/amplicon combination (750 reads per sample) was sufficient to minimize the minor effects of index hopping on the SARS-CoV-2 alignment rate for a given sample, and accurately represent the abundance of SARS-CoV-2-mapping reads in a given samples’ data. Based on the RT-qPCR diagnostics results, samples for which GAPDH was undetectable after 40 cycles of RT-qPCR were removed, and finally 83 patient samples in technical quadruplicates remained.

**Figure 2.**
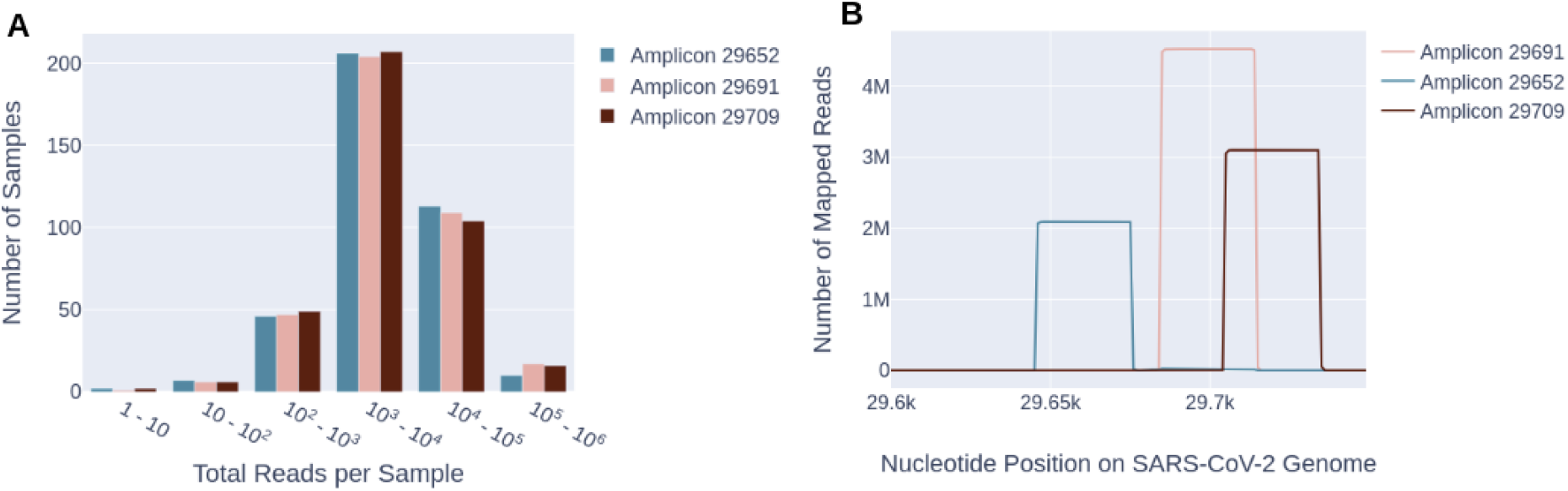
(A) Read distribution in the different wells of the plate. The histogram shows the read numbers obtained in the different wells of the plate for the different amplicons. As the input for the protocols was not normalized, a wide distribution in the number of reads is observed. Wells with <250 reads were discarded for further analysis. **(B) Read alignment to SARS-CoV-2 genome.** The number of reads aligning to the 3’-end of viral genome is shown for the different amplicons. More than 99.9% of reads aligning to SARS-CoV-2 mapped to the locus targeted in the amplicon design.

The alignment of reads to SARS-CoV-2 and GAPDH was performed using Bowtie2 using end-to-end mode. More than 99.9% of mapped reads mapped uniquely to the targeted locus, indicating successful primer design (**Figure 2B**). The global alignment rate of the reads was of 26.96%, 43.47% and 36.69% for amplicons 29652, 29691, and 29709, respectively.

### Diagnostic capability of the assay

Ct values for sets of RT-qPCR technical replicates were normalized and correlated to the alignment rate using the Wilcoxon-Signed-Rank test for matched pairs (p<0.001 for each amplicon; see **Figure 3A**). The per sample alignment rate is shown to be highly correlated with RT-qPCR Ct value (see **Figure 3B**). Samples which returned a Ct value > 25 for SARS-CoV-2 in the diagnostic test, had their alignment rates fall, typically within one standard deviation of the mean alignment rate, for the set of samples diagnosed “negative” with RT-qPCR. This demonstrates that HiDRA-seq is thus far, incapable of identifying with certainty positive samples with a RT-qPCR Ct value > 25. However, there were two exceptions in which the HiDRA-seq system was able to successfully diagnose samples with Ct values > 37. Amplicon 29691 generated the results most consistent with the RT-qPCR diagnosis for Ct values within the range 29 – 40 (see **Figure 3C**).

**Figure 3.**
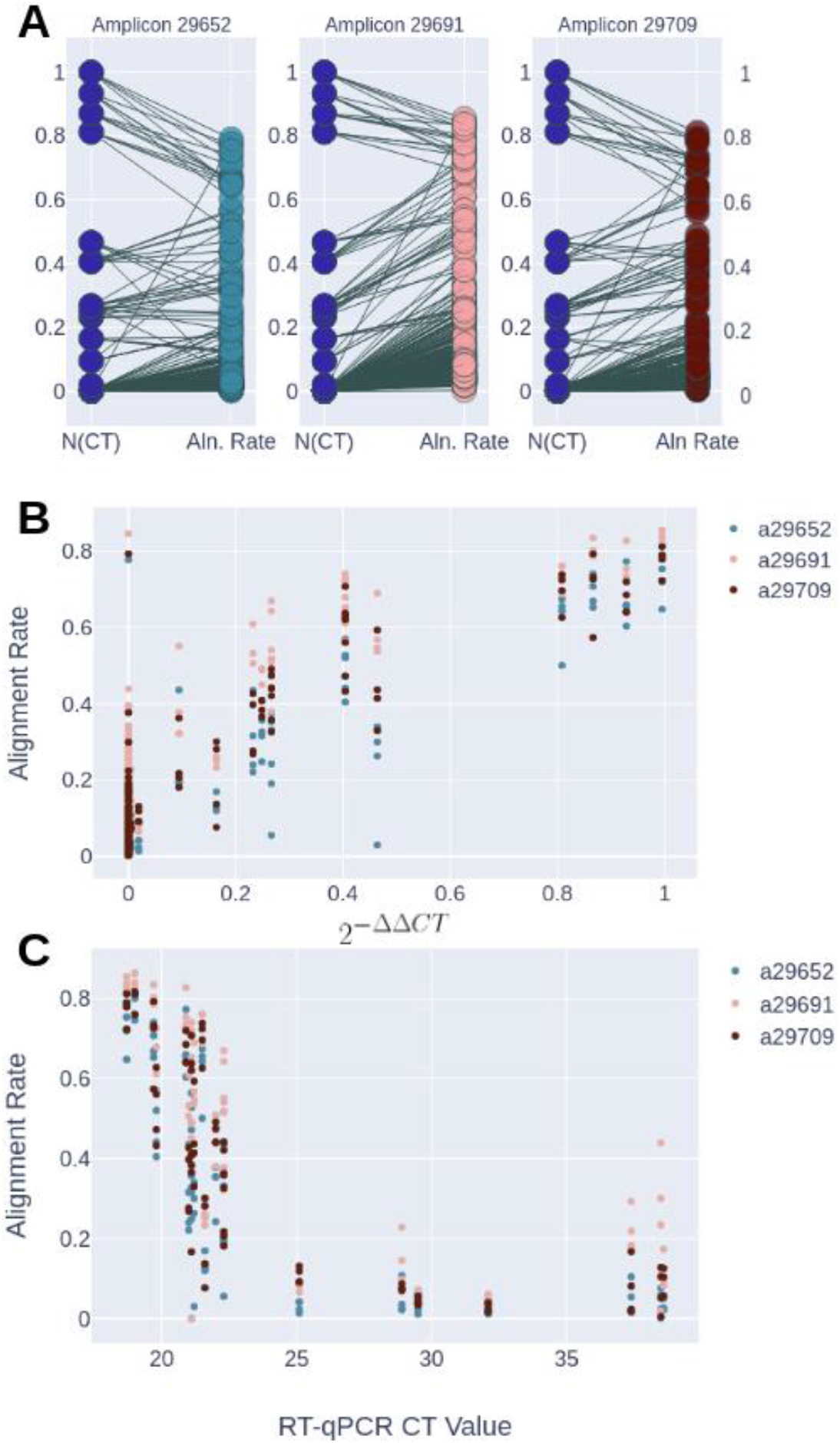
(A) Matched pairs of normalized Ct values and alignment rate. Ct values (y-axis-left) for sets of RT-qPCR technical replicates were normalized 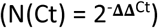 and correlated to the alignment rate (y-axis-right) for the different amplicons using the Wilcoxon-Signed-Rank test for matched pairs (p < 0.001). **(B) Correlation of normalized Ct values and alignment rate.** One point is shown for each sample and amplicon combination, coloured by amplicon type. The normalised Ct value on the x-axis, mapped against its corresponding alignment rate on the y-axis. **(C) Alignment rate vs. raw Ct values for samples diagnosed positive via RT-qPCR.** This figure shows one point for each sample and amplicon combination, and displays only those samples that were diagnosed positive via RT-qPCR. The x-axis shows the Ct value associated with these positive samples, with their corresponding alignment rate shown on the y-axis. This demonstrates a sensitivity threshold for HiDRA-seq in terms of RT-qPCR Ct values (Ct ≈ 25).

We generated potential diagnostic thresholds, based upon the alignment rate (AR) of a sample’s reads to the SARS-CoV-2 genome, by iterating through values on [0,1] in increments of 0.01. We counted for each value, the number of positively diagnosed samples (via RT-qPCR) that had ARs below this value, and the number of negatively diagnosed samples (via RT-qPCR) that had ARs above this value. We then selected intervals for which the number of false diagnoses were minimized as a basis for a theoretical diagnostic threshold. Potential diagnostic thresholds were set individually for each amplicon as follows (see **Figure 4**): Amplicon 29652: AR ∊ [0.02, 0.11]; Amplicon 29691: AR ∊ [0.05, 0.24]; Amplicon 29709: AR ∊ [0.04, 0.18]. On the intervals of AR values for which the number of false diagnoses was minimized, amplicons 29652 and 29709 mis-diagnosed 8 patients and Amplicon 29692 mis-diagnosed 7 patients. Furthermore, amplicons 29652, 29709 and 29692 had a successful diagnostic rate with respect to RT-qPCR diagnosis of 90.4%, 90.4% and 91.6%, respectively.

**Figure 4.**
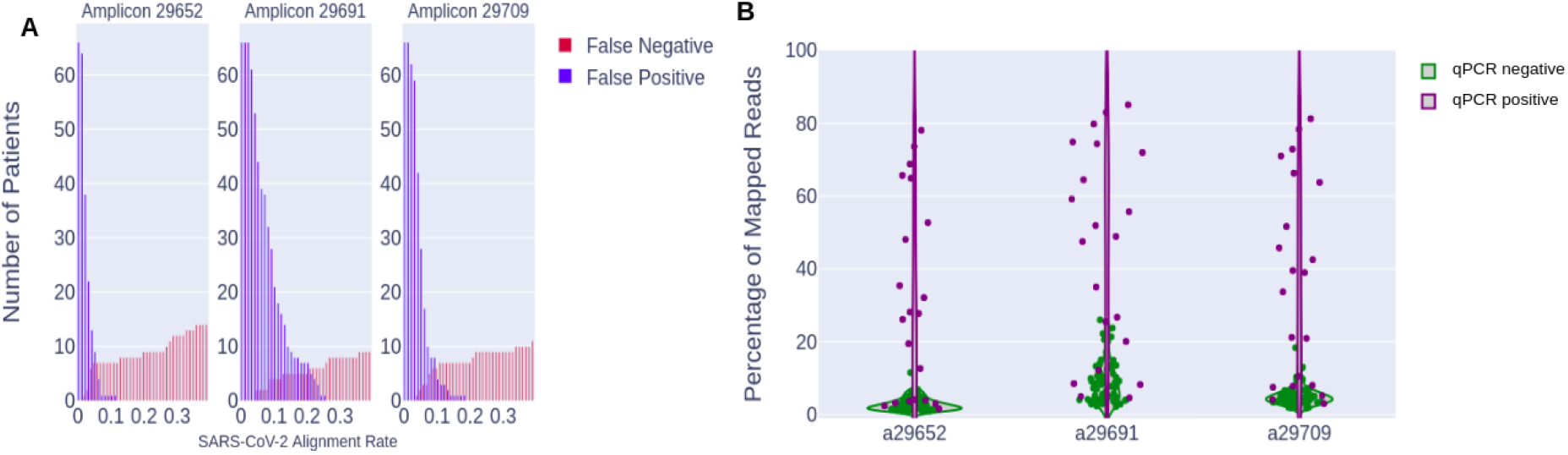
(A) Histogram of false diagnoses for potential diagnostic thresholds: For each amplicon used, the number of potential mischaracterized diagnoses (false-positives in blue or false-negatives in red), in numbers of patients (y-axis), are shown for a given alignment rate threshold (x-axis). Original diagnoses are given via RT-qPCR. Amplicon 29691 is shown to have the fewest number of mis-characterized diagnoses (7) when said diagnoses are minimized, and Amplicon 29652 is shown to have the shortest span of intersection between both distributions (0.9). **(B) Distributions of alignment rates for positive and negative patient quadruplicates, by amplicon:** For each amplicon, the mean percentage (across four replicates) of reads that aligned to the SARS-CoV-2 genome (y-axis) is shown as a point, with patients diagnosed positive via RT-qPCR shown in purple and patients diagnosed negative via RT-qPCR shown in green. The distribution of alignment rates for patients diagnosed positive with RT-qPCR is shown to be much wider (spanning roughly 87 percentiles on average) than those diagnosed negative (spanning roughly 14 percentiles on average).

## Discussion

The surge in cases of SARS-CoV-2 infections around the world has created an urgent need for accurate and fast diagnostic testing solutions. Despite the recent increase in available SARS-CoV-2 diagnostic testing kits in the past few months, the majority of tests still rely on real-time quantitative PCR (RT-qPCR). Whilst RT-qPCR reactions are generally very sensitive (*i.e.* able to detect true positive cases) and specific, the technology has inherent limitations with regard to large scale population screening, which has become increasingly important during this pandemic. Additionally, RT-qPCR does not provide any genotypic information regarding a patient’s infection beyond the causal organism. Another advantage of NGS over RT-qPCR is that NGS provides a direct and functional measurement SARS-CoV-2. RT-qPCR generates florescent measurements for individual plate wells, indirectly quantifying the presence of genetic material in relative concentrations. Alternatively, NGS generates thousands of reads, directly measuring the specific sequences present in a sample. The sequence information generated could provide insight into the specific infecting isolate and aid in tracing transmission within communities.

HiDRA-seq, built on Next Generation Sequencing technology, has the ability to multiplex thousands of barcoded patient samples, significantly increasing current testing capacity. Our method is designed to be partially performed on a small automated liquid handling machine, so that a single person is able process more than 2,000 RNA samples per day with ease. This results in an overall shorter diagnostic turnaround time, with library preparation and sequencing data obtained in as little as 1.5 days. Our system does not match the speed of RT-qPCR for individual samples. However, it outperforms standard diagnostics methods in scale, allowing one to process hundreds of thousands of samples per week if extended and automated at scale. The faster a test can be administered, the sooner the results can be received, and the quicker measures can be put in place to mitigate further spread or to evaluate the impact of loosened containment measures. This system also minimizes the errors in sample handling by fast-tracking sample preparation, an integral part of the workflow. The achievement of consistent data across samples verifies the reproducibility and reliability of our system. Additionally, the miniaturization of our reactions results in a much more affordable solution compared to other methodologies available in the market. Our estimated price for sample screening from extracted RNA to diagnostic result with our approach is 2 CHF/EUR/USD.

By comparing our results to the RT-qPCR-based clinical diagnostic test, we show that the mapping rate is a strong predictor of Ct values (p<0.001). Our method is sensitive, as we have been able to correctly recall positive samples in >90% of the cases, which is comparable to other NGS-based methods use for virus detection in clinical samples^7^. The samples that escaped detection are characterized by having high Ct values from the diagnostic RT-qPCR (Ct>25). In parallel to HiDRA-seq, we prepared libraries using the highly sensitive Smart-seq2 protocol^8^, which captures all poly-adenylated RNA in the sample, and we were similarly unable to detect a significant number of virus-derived reads. The sensitivity of HiDRA-seq could be improved by slightly increasing the number of PCR cycles used in the amplicon generation and using UMI correction for detection of PCR duplicates to increase the quantitative accuracy of the method. Although we were able to successfully diagnose the majority of samples using 50 bp reads, the designed primers allowed us to access 71 bp of a highly variable region (following Amplicon 29709) by generating reads of 150 bp, raising the possibility of using this method for basic phylogenetic and epidemiological studies of isolates differing at this genomic position.

HiDRA-seq will be optimized to enable direct lysis from saliva collected by gargling. Given that the mcSCRB-seq^4^ method, from which this protocol derives, is designed to work with direct lysis from single cells, this approach will be adapted for HiDRA-seq, since RNA extraction is one of the biggest bottle-necks for large scale testing. The current sensitivity of our method makes it compatible with direct testing from saliva, as suggested in a recent publication^9^.

Our method uses barcoded oligo-dT primers to generate cDNA in the reverse transcription and this feature leaves the door open to expanding the amplicon panel. To achieve this, additional PCR primers could be added to generate amplicons that are specific for other human coronaviruses (hCoVs) or other respiratory virus that produce polyadenylated transcripts. Such viruses include influenza viruses (IAVs), respiratory syncytial viruses (RSVs), parainfluenza viruses (PIVs) or human metapneumoviruses (MPVs), which would create a multi-viral identification test at almost no extra cost. Potentially, this approach could also be implemented to specifically detect the expression of virtually any human mRNA identified as a biomarker for estimating disease susceptibility and progression, or for designing host group-specific COVID-19 treatment regimens. This is especially relevant in a clinical research setup, in which the importance of a test that could both identify an infection and give information on how to best treat that infection cannot be overstated.

This method has been designed with the practical necessities of large scale, affordable, adaptable and rapid testing in mind. To these ends, we have developed a first version of a method that reuses relatively common sets of barcoding primers available in NGS facilities, can scale effectively, does not involve exotic reagents and relies on NGS to multiplex samples for both cost and time savings, allowing any well-equipped sequencing lab in the world to quickly begin testing.

## Materials and Methods

### Primer sequences

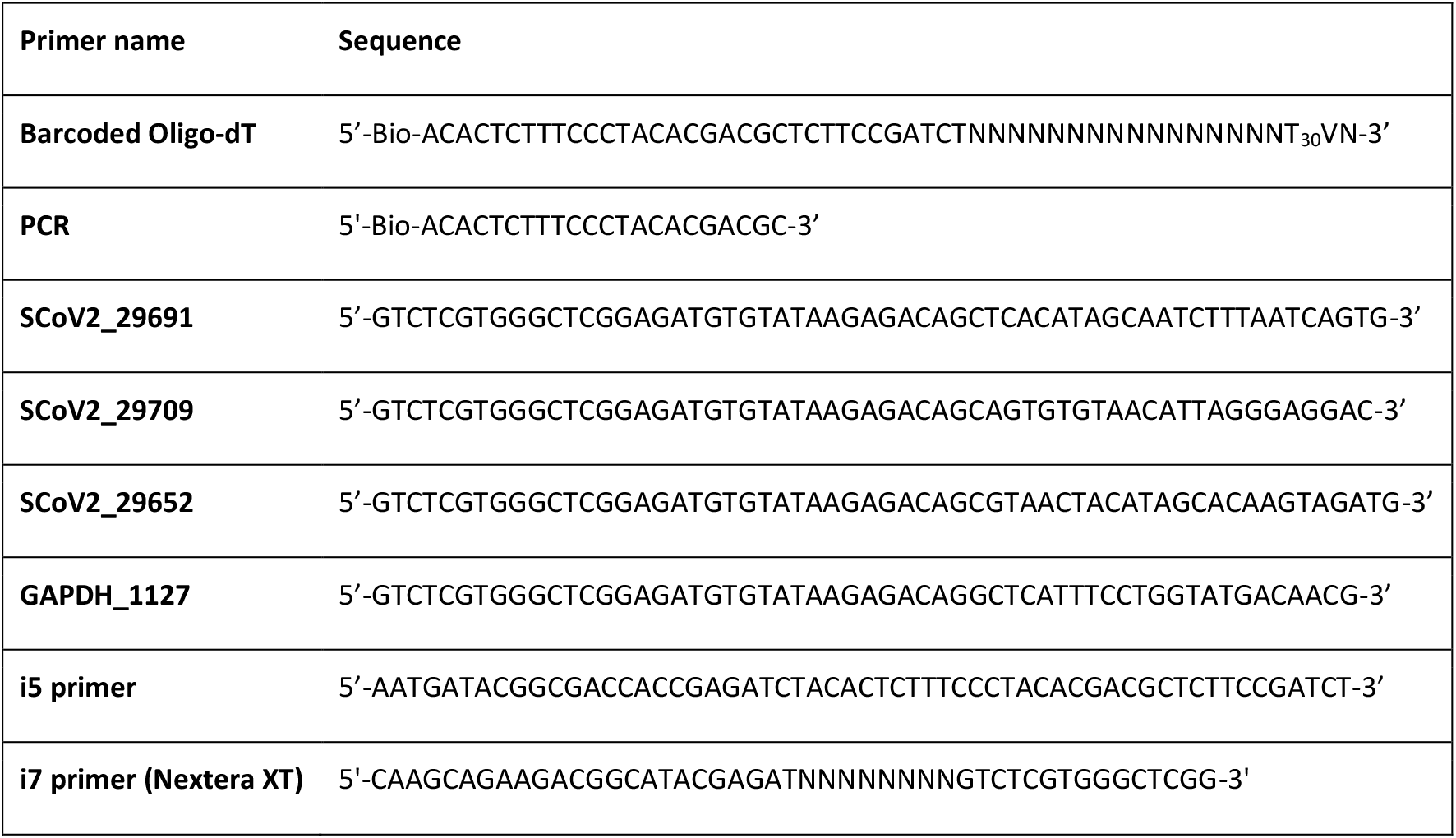

### Reverse transcription with barcoded oligo-dT

This part of the protocol is based on the reverse transcription strategy used in mcSCRB-seq (Bagnoli et al., 2018). A 384-well barcoding plate was prepared using a Mosquito HV liquid handling robot (SPT Labtech), each of the wells containing 0.5 μl of lysis solution (0.2% Triton X-100 [Roche], 0.8 units of RNasin Plus [Promega], 4 mM dNTPs [Promega] and 1 μM of barcoded oligo-dT primers [E3V6NEXT, IDT]). The plate was divided into four quadrants (91 samples each) with four identical copies of a 96-well plate containing RNA extracted from anonymous patients’ swaps (kindly provided by Dr. Michael Huber and Dr. Jürg Böni, Institute of Medical Virology, UZH, Switzerland) by transferring 0.5 μl of RNA to the corresponding well positions. The plate was heated at 65° C for 5 minutes and transferred to ice prior to the addition of 1 ul of 2X Reverse transcription mix (15% PEG 8000 [Sigma Aldrich], 2X Maxima RT buffer [Thermo Fischer] and 4 units of Maxima H Minus RT [Thermo Fisher]). cDNA synthesis was performed for 15 min at 50°C followed by 5 min at 85° C for inactivation. The Mosquito HV liquid handling robot (SPT Labtech) was used to pool the whole 384-well plate containing the barcoded cDNA into a single 2 ml DNA LoBind tubes (Eppendorf) and cleaned up using Sera-Mag Select beads (GE Healthcare) with a ratio 1:0.8 (pooled cDNA:beads). Purified cDNA was eluted in 17 μl and residual primers digested with Exonuclease I (Thermo Fisher) for 20 min at 37 °C.

### SARS-CoV-2-specific amplicon generation

Three different SARS-CoV-2-specific primers (SCoV2_29691, SCoV2_29709, SCoV2_29652) were used in individual PCR reactions, in combination with a human GAPDH-specific primer (GAPDH_1127) and a PCR primer binding to the barcoding sequence, to generate three virus specific amplicons. These primers also contain the adaptor sequences needed to incorporate the Illumina compatible flow cell binding sequences in a subsequent PCR reaction. Briefly, each PCR reaction was assembled by combining 5 ul of the pooled barcoded cDNA with 20 μl of PCR master mix (1.25X KAPA HiFi HotStart ReadyMix [Roche], 0.375 uM of SARS-CoV-2- and GAPDH-specific primers [Microsynth AG] and 0.375 uM of the PCR primer [Microsynth AG]). PCR was performed using the following program: 3 min at 98 °C for initial denaturation followed by 25 cycles of 20 sec at 98 °C, 15 sec at 60 °C, 15 sec at 72 °C. Final elongation was performed for 5 min at 72 °C. Once the PCR was concluded, the amplicons were cleaned up using Sera-Mag Select beads (GE Healthcare) with a ratio 1:0.8 (DNA:beads). The size and concentration of the amplicons was analyzed in a 4200 TapeStation System (Agilent) using a D1000 ScreenTape.

### Library generation and plate barcoding incorporation

A final PCR was performed from the amplicon to generate libraries compatible with Illumina sequencer. In parallel, three different barcodes were assigned for the aforementioned amplicons in order to combine them in a single sequencing run. This strategy can be used to combine different plates with patient samples, allowing one to easily increase the throughput of the protocol. Each PCR reaction was assembled by combining, in individual tubes, 5 ul of the three amplicons with 20 μl of PCR master mix containing 1.25X KAPA HiFi HotStart ReadyMix (Roche), 0.375 uM of the i5 primer (Microsynth AG) and 0.375 uM of the barcoded i7 primer (Illumina). PCR was performed using the following program: 3 min at 98 °C for initial denaturation followed by 5 cycles of 20 sec at 98 °C, 15 sec at 55 °C, 15 sec at 72 °C. A final elongation was performed for 5 min at 72 °C. Once the PCR was completed, the libraries were cleaned up using Sera-Mag Select beads (GE Healthcare) with a ratio 1:0.8 (DNA:beads). The size and concentration of the libraries were analyzed in a 4200 TapeStation System (Agilent) using a D1000 ScreenTape.

### Sequencing

The three libraries generated from three different viral amplicons, all in combination with a GAPDH-derived library, were paired-end sequenced together in both an Illumina MiniSeq (R1=16 bp, i7=8 bp, R2=50 bp) and NovaSeq6000 sequencers (data not shown, R1=16 bp, i7=8 bp, R2=150 bp). PhiX was added to account for 25% of the total library, to increase library diversity and subsequently, sequencing performance. The 6 first bases in read one were used to demultiplex the reads generated from the different patient. The 8 bases in the barcode allowed, in this case, to discriminate the different amplicons but it could alternatively be used to multiplex several patient plates for sequencing. The 50/150 bases in read 2 were used to distinguish between positive and negative patients by mapping reads against the SARS-CoV-2 genome.

### Bioinformatic analysis

Reads were segregated by patient, amplicon and plate. The demultiplexed reads were then processed using the standard tools displayed below. A consensus sequence for SARS-CoV-2 genome was generated from the set of published genomes on NCBI using the Bio.Align Python 3 package (data not shown). Bowtie2^10^, samtools^11,12^ and pysam were used to generate pileup columns and calculate alignment rates for a subset of our samples, on all loci in the SARS-CoV-2 transcriptome to verify the locus-specificity of our primers. After verifying the quality of our reads, Bowtie2 was used to map all of our samples against the SARS-CoV-2 genomic region of interest (from 29600 bp to 29900 bp, inclusive), and GAPDH. Alignment rates were calculated as a ratio of total reads for a given sample, to the number of reads aligning to our region of interest on the SARS-CoV-2 genome.

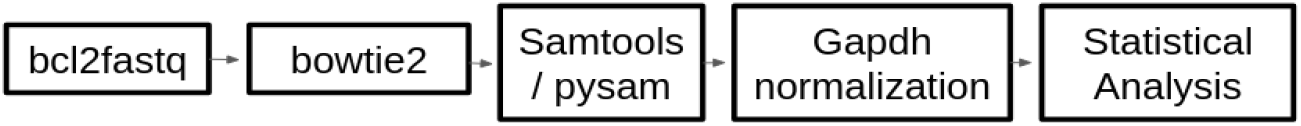

Samples for which the initial diagnostic PCR failed (*i.e.* GAPDH was undetectable after 40 cycles of PCR) were filtered from the dataset. Additionally, samples for which we recovered less than 250 sequencing reads per amplicon were filtered out of the dataset. After filtering had been applied, our dataset contained 83 sets of patient quadruplicates. Undetected Ct values of SARS-CoV-2 were imputed and values were then normalized to GAPDH using the Livak and Schmittgen method^13^. The Wilcoxon Signed-Rank Test was applied to the set of patient replicates, comparing the SARS-CoV-2 alignment rate and the normalized Ct value recovered from RT-qPCR (p < 0.0001 for each amplicon).

## Author contributions

CA and RS conceived the study. EY, GW, SK, LP and CA designed the protocol, prepared the libraries and sequenced them. Sequencing data was processed and analysed by GW and LO. EY, GW, SK, LO, LP, TS, MDM, CA and RS wrote the manuscript.

## Acknowledgments

We thank Dr. Michael Huber and Dr. Jürg Böni (Institute of Medical Virology, University of Zurich, Switzerland) for providing the RNA extracted from patients’ samples used for the implementation of the protocol. We are grateful to Prof. Andreas Moor (D-BSSE, ETH Basel, Switzerland) for providing the set of 384 barcoded oligo-dT primers used in this protocol.

## Competing interests

The authors declare no competing interests.

## Data availability

Sequencing data generated here are available at ENA under accession PRJEB38511.

## Notes

### Competing Interest Statement

The authors have declared no competing interest.

